# The Subcortical Atlas of the Marmoset (“SAM”) monkey based on high-resolution MRI and histology

**DOI:** 10.1101/2024.01.06.574429

**Authors:** Kadharbatcha S Saleem, Alexandru V Avram, Daniel Glen, Vincent Schram, Peter J Basser

## Abstract

A comprehensive three-dimensional digital brain atlas of cortical and subcortical regions based on MRI and histology has a broad array of applications for anatomical, functional, and clinical studies. We first generated a **S**ubcortical **A**tlas of the **M**armoset, called the “SAM,” from 251 delineated subcortical regions (e.g., thalamic subregions, etc.) derived from the high-resolution MAP-MRI, T2W, and MTR images *ex vivo*. We then confirmed the location and borders of these segmented regions in MRI data using matched histological sections with multiple stains obtained from the same specimen. Finally, we estimated and confirmed the atlas-based areal boundaries of subcortical regions by registering this *ex vivo* atlas template to *in vivo* T1- or T2W MRI datasets of different age groups (single vs. multisubject population-based marmoset control adults) using a novel pipeline developed within AFNI. Tracing and validating these important deep brain structures in 3D improves neurosurgical planning, anatomical tract tracer injections, navigation of deep brain stimulation probes, fMRI and brain connectivity studies, and our understanding of brain structure-function relationships. This new *ex vivo* template and atlas are available as volumes in standard NIFTI and GIFTI file formats and are intended for use as a reference standard for marmoset brain research.

## Introduction

The basal ganglia, thalamus, hypothalamus, brainstem nuclei, and amygdala are subcortical regions that regulate sensorimotor, cognitive, limbic, and autonomic (sympathetic and parasympathetic) functions (Lanciego et al. 2012; Ferrazzoli et al. 2018; Sklerov et al. 2019; Aggleton and O’Mara 2022). A comprehensive 3D digital template atlas of these subcortical regions in non-human primates (NHP) based on MRI and histology is of great value in anatomical, functional, and clinical studies. In particular, a 3D atlas can be registered to a NHP brain MRI volume to best determine the regions of interest for anatomical tracer injections for connectome studies, the areal location of observed fMRI responses, and the potential targets for deep brain stimulation (DBS) in NHP models of neurological disorders. In this study, we developed a **S**ubcortical **A**tlas of the **M**armoset (**SAM**) monkey using an ultra-high-resolution Mean Apparent Propagator (MAP)-MRI, T2W, and MTR images combined with a corresponding set of histological images from the same marmoset brain.

MAP-MRI (Ozarslan et al. 2013) provides a comprehensive and clinically feasible (Avram et al. 2016) assessment of water diffusion in tissues. In each voxel, the MAP-MRI explicitly measures the probability density function of the net 3D microscopic displacements of diffusing water molecules, also known as diffusion propagators. MAP-MRI subsumes and generalizes other diffusion MRI (dMRI) signal representations (Avram et al. 2017) and quantifies average diffusion properties in isotropic and anisotropic tissues with arbitrary microstructure and architecture thoroughly and accurately with multiple microstructural markers. Besides the DTI-derived fractional anisotropy (FA), axial, radial, and mean diffusivities (AD, RD, and MD, respectively), direction-encoded color (DEC) maps (Pajevic and Pierpaoli 1999), MAP-MRI yields a family of new microstructural parameters that capture more subtle features of diffusion propagators, such as the zero-displacement probabilities, the non-gaussianity index (NG), and the propagator anisotropy (PA) (Ozarslan *et al*. 2013; Avram, Barnett, et al. 2014; Avram *et al*. 2017; Avram, Bernstein, et al. 2018), as well as MAP-derived orientation distribution functions (ODFs). Taken together, the DTI/MAP parameters provide a more sensitive and specific microstructural assessment compared to conventional dMRI (Hutchinson et al. 2018) and structural (e.g., T1W and T2W) scans. They have proven remarkably effective for detailed anatomical segmentation of the cortical (Avram, Saleem, Komlosh, et al. 2022) and subcortical structures (Saleem et al. 2021; Saleem et al. 2023).

Many studies have segmented the subcortical structures and provided 3D atlases in humans using high-resolution *in vivo* MRI with multiple image contrasts or *ex vivo* spin echo (SE) T2W MRI with histological stains (Rijkers et al. 2007; Abosch et al. 2010; Lenglet et al. 2012; Deistung, Schafer, Schweser, Biedermann, Gullmar, et al. 2013; Deistung, Schafer, Schweser, Biedermann, Turner, et al. 2013; Keuken et al. 2014; Ewert et al. 2018; Pauli et al. 2018; Hoch, Bruno, Faustin, Cruz, Crandall, et al. 2019; Hoch, Bruno, Faustin, Cruz, Mogilner, et al. 2019). In contrast, only a limited number of studies have provided detailed maps of subcortical regions or created a 3D digital subcortical atlas template using combined MRI and histology in the marmoset. 1) Newman and colleagues described a subcortical atlas of a marmoset monkey (Newman et al. 2009). However, it is a non-digital version based on the series of individual photographs of Nissl sections with closely matched MR images obtained from a different marmoset brain. A similar type of histology-based atlases in book form has been published with or without MRI (Stephan et al. 1980; Palazzi and Bordier 2008; Yuasa et al. 2010; Hardman and Ashwell 2012; Paxinos et al. 2012; Iriki et al. 2018). 2) The Brain/Minds digital-marmoset brain atlas has provided the 3D segmentation of major cortical and subcortical regions using a standard structural T1/T2W MRI, coregistered with only Nissl-stained histology data to identify the region-of-interest (Hashikawa et al. 2015; Woodward et al. 2018). 3) Liu and colleagues (Liu et al. 2018) constructed a 3D digital atlas of the marmoset brain based on *ex vivo* MTR, T2W, and diffusion MRI. This study provided the segmentation of 54 cortical, but few subcortical (n=16) regions in their atlas, and the area labels were derived from a different subject (Paxinos *et al*. 2012). 4) A recent study generated a histology-based probabilistic atlas of the marmoset brain called the Nencki-Monash template, but it was focused only on the cytoarchitectonic parcellation of cortical areas (Majka et al. 2020; Majka et al. 2021). Similar to the marmoset, only a few studies have produced the detailed mapping of subcortical regions in the macaque monkey using combined MRI and histology (for more details, see the introduction section in Saleem et al., 2021).

Many of the subcortical nuclei and their subregions are challenging to identify and delineate in conventional MRI due to their small size, buried location, and often subtle contrast compared to neighboring regions. As shown in our previous study in the macaque (Saleem *et al*. 2021) and marmoset (Saleem *et al*. 2023) monkeys, combining volumes of different MRI markers acquired with high-spatial resolution (100-200 μm), aided by whole-brain histological information derived from the same brain specimen, is key to delineating nuclei and fiber tracts in deep brain structures, including substructures and laminae, e.g., in the thalamus and basal ganglia. Thus, the segmentation and validation of brain areas based on MRI-histology correlations are crucial for the construction of accurate 3D digital template atlases in NHPs.

In this study, we combined MAP-MRI, T2W, and MTR data with the matched high-resolution images of histology sections with multiple stains derived from the same *ex vivo* marmoset brain specimen to segment 251 subcortical regions and associated white matter pathways. This integrated multimodal approach yields a more objective and reproducible delineation of gray matter nuclei and their boundaries in the deep brain structures, including the basal ganglia, thalamus, hypothalamus, limbic region (amygdala), basal forebrain, and the rostrocaudal extent of the brainstem (midbrain, pons, and medulla). We derived the 3D Subcortical Atlas of the Marmoset brain (SAM) from these segmented deep brain regions and registered this volume to a range of *in vivo* T1- and T2W standard MRI volumes from control subjects in different age groups to illustrate the application of the atlas to in vivo studies. This newly derived *ex vivo* 3D digital atlas is intended to provide a practical standard template for neuroanatomical, functional (fMRI), clinical, and connectional imaging studies involving subcortical targets in marmoset monkeys.

## Materials and Methods

### Perfusion fixation

One adult male marmoset monkey (*Callithrix jacchus*), weighing 340 gm, was used in this study. This animal was previously involved in the transgenic studies at the NIMH/NIH transgenic core, and we received the perfusion-fixed brain specimen of this animal from this core for our *ex vivo* MRI and histological studies. All procedures adhered to the Guide for the Care and Use of Laboratory Animals (National Research Council) and were carried out under a protocol approved by the Institutional Animal Care and Use Committee of the National Institute of Mental Health (NIMH) and the National Institutes of Health (NIH). The animal was deeply anesthetized with sodium pentobarbital and perfused transcardially with 0.5 liters of heparinized saline, followed by 2 liters of 4% paraformaldehyde, both in 0.1M phosphate buffer (pH 7.4). After perfusion, the brain was removed from the cranium, photographed, and post-fixed for 24 h in the same buffered paraformaldehyde solution and then transferred into 0.1 M phosphate-buffered saline (PBS) with sodium azide before MRI scanning.

### Ex vivo MRI

#### Data acquisition

The MRI data acquisition and processing are described in detail in our previous study (Saleem *et al*. 2023). In brief, we positioned the fixed marmoset brain specimen in a 3D printed brain mold and then inside a custom 30 mm diameter cylindrical container. We then filled the 3D mold and container with fomblin and prepared the sample for MR imaging using a Bruker 7T/300 mm horizontal MRI scanner with a 30 mm inner diameter quadrature millipede coil (ExtendMR; http://www.extendmr.com/).

We acquired MAP-MRI data using a diffusion spin-echo (SE) 3D echo-planar imaging (EPI) sequence using the following imaging parameters: 150 µm isotropic resolution, 4.32×2.76×2.76 cm^3^ field-of-view (FOV), 288×184×184 imaging matrix, ten segments per kz-plane, and 1.33 partial Fourier acceleration. We acquired 256 diffusion-weighted images (DWIs) with diffusion gradient pulse durations and separations of δ=6 ms and Δ=28 ms, respectively, a 48 ms echo time (TE) and a 650 ms repetition time (TR). MAP-DWIs were acquired using multiple b-value shells: 100, 500, 1000, 1500, 2500, 3500, 4500, 5250, 7000, 8500, 10000 s/mm^2^ with uniformly sampled gradient directions (Koay et al. 2012; Avram, Sarlls, et al. 2018) on each shell (7, 10, 12, 15, 21, 22, 27, 31, 35, 36, and 40, respectively). We also obtained a magnetization transfer (MT) prepared scan using a 3D gradient echo sequence with 150 µm isotropic resolution (288×184×184 imaging matrix, 4.32×2.76×2.76 cm FOV with a 15° excitation flip angle, TE/TR=3.7/37ms, two averages, MT saturation pulse with 2 kHz offset, 12.5 ms Gaussian pulse with 6.74 µT peak amplitude, and 540° flip angle). The total duration of the MAP-MRI scan was 85 h and 20 min, and the MT scan was 11 h and 8 min.

#### Data processing

From the scans acquired with and without MT, we computed the MT ratio (MTR) volume with good GM/WM tissue contrast to serve as a structural template for subsequent DWI distortion correction and co-registration. We processed the MAP-MRI dataset with the comprehensive Tortoise pipeline (Pierpaoli et al. 2010) that performs denoising, corrects for Gibbs ringing, gradient eddy currents, intra-scan drift, and EPI distortions, and registers all volumes to the MTR scan. We estimated diffusion propagators by fitting the data in each voxel with a MAP series truncated at order four and computed DTI (MD, AD, RD-mean, axial, and radial diffusivities, respectively); FA-fractional anisotropy; CL, CP, CS-linear, planar, and spherical anisotropy coefficients, respectively (Westin et al. 2002) and MAP-MRI tissue parameters (RTOP-return-to-origin probability, RTAP-return-to-axis probability, RTPP-return-to-plane probability, PA-propagator anisotropy, and NG-non-Gaussianity), along with the corresponding baseline (T2W) volume. Furthermore, we estimated and visualized fiber orientation distribution functions (FODs) using MRtrix 3.0.1 (Tournier et al. 2012). For more details, see our previous study (Saleem *et al*. 2023).

#### Histological processing and data analysis

All the histological procedures (section cutting and staining) are described in detail in Saleem et al. (2023). Following MRI acquisition, we prepared the whole brain specimen as one tissue block for histological processing with five different stains. Serial frozen coronal sections (50 µm thick) were cut on a sliding microtome through the entire brain, including the cerebrum, brainstem, and cerebellum. We collected a total of 670 sections from this brain, sorted them into five parallel series (134 sections per set with 250 μm spacing between adjacent sections), and stained them sequentially for parvalbumin (PV), neurofilament (SMI 32), acetylcholinesterase (AchE), Cresyl violet (CV), and choline acetyltransferase (ChAT), respectively (Fig. 1). We stained all the sections in this animal and also collected block-face images of a frozen tissue block at every 250 μm interval. The commercially available antibodies used for SMI-32, PV, and ChAT staining are indicated in the “*Key Resources Table*.”

**Fig. 1.**
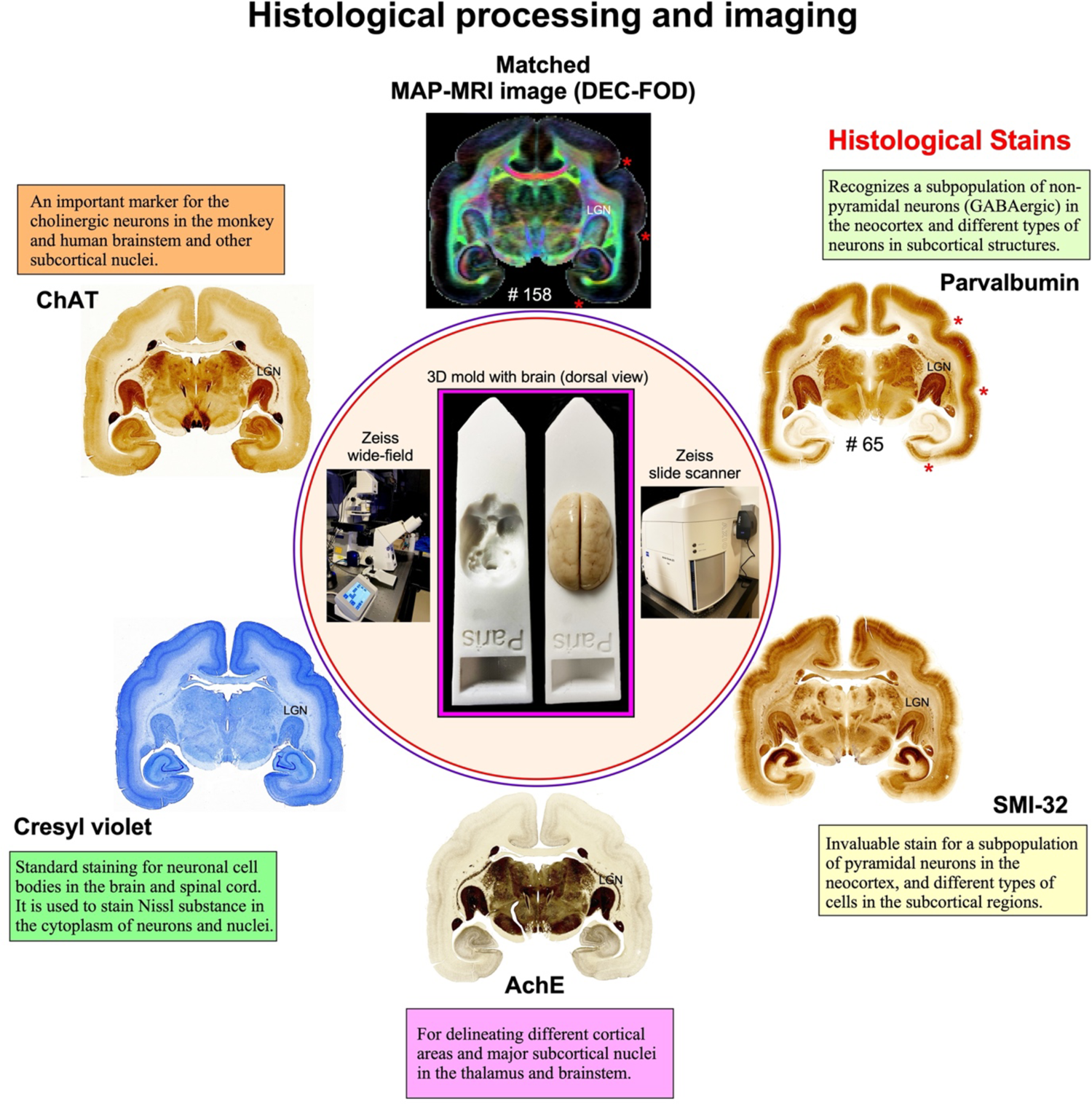
Histological staining and high-resolution imaging. Frozen sections were cut coronally from the frontal cortex to the occipital cortex at 50 μm thickness on a sliding microtome. In total, 670 sections were collected, and all the sections were processed with different cell bodies and fiber stains. This example shows an adjacent series of five sections at the level of the anterior temporal cortex stained with parvalbumin, SMI-32, AchE, Cresyl violet, and ChAT, respectively. The unique characteristics of each stain are described in the “*histological processing and data analysi*s” section. We obtained 134 stained sections in each series, and the interval between two adjacent sections in each series is 250 μm. The high-resolution images of stained sections were captured using a Zeiss wide-field microscope and a Zeiss Axioscan Z1 high-resolution slide scanner (inset). These histology images were then aligned manually with the corresponding MAP-MRI (top) and other MRI parameters of the same specimen to allow visualization and delineation of subcortical structures in a specific region of interest. In both MRI and histology sections, note the correspondence of sulci (red stars), gyri, and deep brain structures (e.g., LGN-lateral geniculate nucleus). #65 refers to the section number in each set/series of stained sections, and #158 indicates the matched MRI slice number in 3D volume. The inset shows the 3D brain mold with and without the brain specimen for MR imaging.

#### Histology data analysis

High-resolution images of all stained sections were captured using a Zeiss wide-field microscope and Zeiss high-resolution Axioscan-Z1 slide scanner at 5X objective, and these digital images were adjusted for brightness and contrast using Adobe Photoshop (v24.2). These images were then aligned manually with the corresponding images of DTI/MAP parameters along with the estimated T2W (i.e., non-diffusion weighted) and the MTR images to allow visualization and delineation of subcortical structures in specific region-of-interest (ROIs) (e.g., Figs. 3, 4). Some structures, like striatum and pallidum, were demarcated by juxtaposing matched MRI and histology sections, but for others (e.g., amygdala), we used a different approach to outline their subregions as described in our previous study (Saleem *et al*. 2021). In brief, we first superimposed a histology section onto the matched MRI slice and then manually rotated and proportionally scaled it to match the boundaries of the different nuclei and their subregions on the MRI using the transparency function in Canvas X Draw software 7.0.3. Finally, the borders of the subregions were manually traced on the histology sections and translated onto the superimposed underlying MR images using the polygon-drawing tools and smooth and grouping functions in this software (Saleem *et al*. 2021; Saleem *et al*. 2023).

The histological sections were matched well with the MRIs in this study. It was not required to resample the MRI volume to achieve good alignment with the histological images in the current study. The alignment of these images was only possible by carefully orienting the entire brain specimen with reference to sagittal MR images from this specimen. These steps were done on the microtome stage before attaching and freezing the brain specimen with dry ice. These steps enabled us to match the sulci, gyri, and region of interest (ROI) in deep brain structures in both MRI and histological sections, as shown in Figure 1 and the result section below (Figs. 3, 4).

#### Delineation of subcortical regions and generation of standard “SAM” (ex-vivo) atlas

Subregions (ROIs) of the basal ganglia, thalamus, hypothalamus, amygdala, brainstem, and other deep brain areas and selected fiber bundles were manually segmented through a series of 146, 150 μm thick coronal sections in PA/DEC, T2W, or other MRI parameters using ITK-SNAP (v4.0) (Yushkevich et al. 2006). The spatial extent and borders of each segmented region in MRI were confirmed with the matched high-resolution histology images obtained from multiple stained sections (Fig. 2). The regions were drawn manually only on the left hemisphere. In order to define a good axis of symmetry, the MRI dataset was rotated about the anterior-posterior axis by a small angle of 0.75 degrees in AFNI’s Nudge plugin. The dataset coordinates were set to be aligned to an AC-PC (Anterior Commissure – Posterior Commissure) axis with the center of the Anterior Commissure at the (0,0,0) XYZ coordinate.

**Fig. 2.**
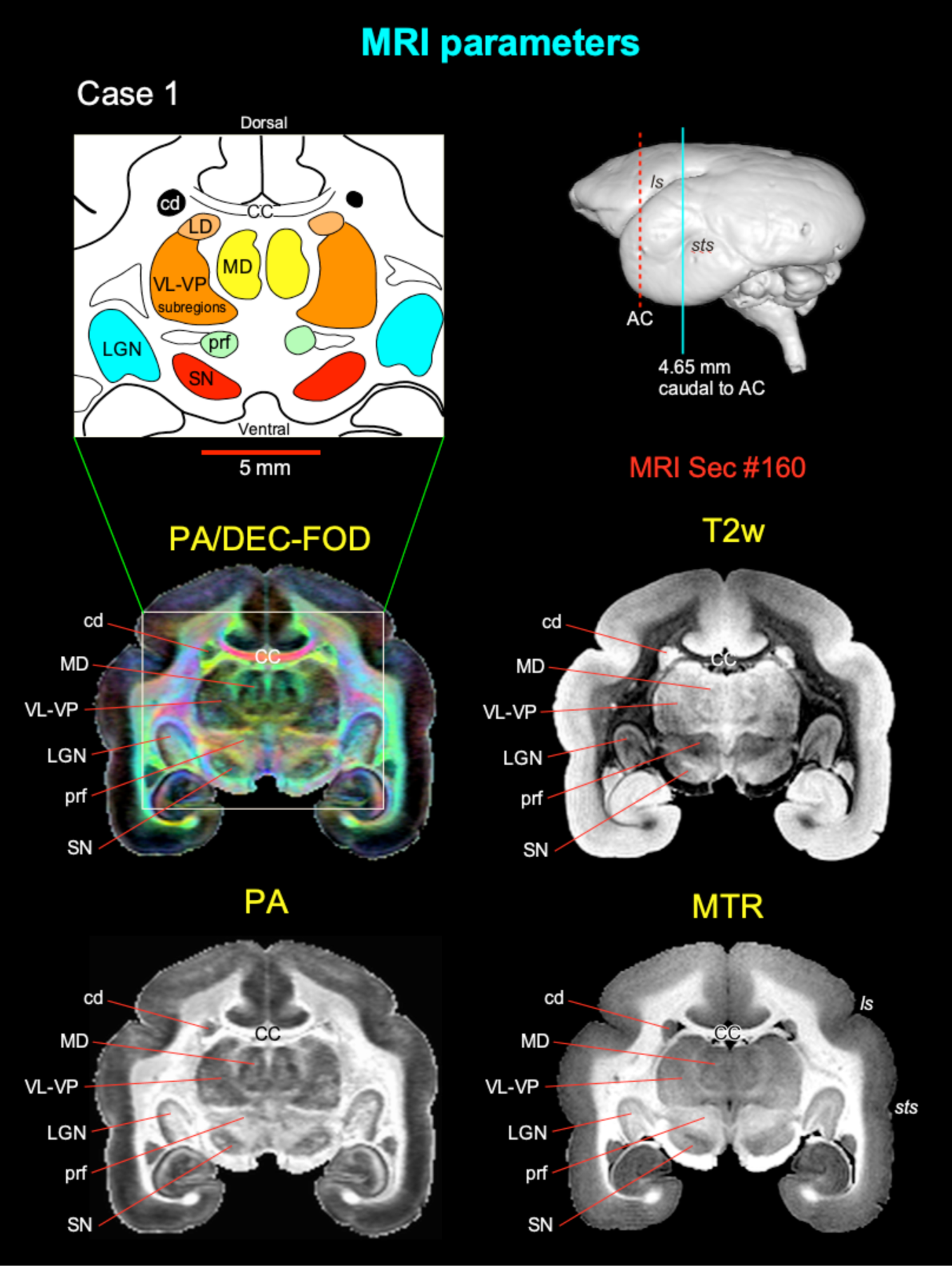
Subcortical regions in different MRI parameters. Matched coronal MR images from two of the eight MAP-MRI parameters, T2-weighted (T2W), and the magnetization transfer ratio (MTR) images show selected subcortical regions: thalamic subregions (MD, LD, VL, VP, LGN), basal ganglia subregions (SN, cd), and a prerubral region (prf) anterior to the red nucleus. These areas are also illustrated in the corresponding drawing from the PA/DEC-FOD image on the top left (white box/inset). For mapping and detailed description of the thalamic and other subcortical regions and other MAP-MRI parameters, see the result section in Saleem et al. (2023). This MRI slice is located at the level of the rostral temporal cortex and 4.65 mm caudal to the anterior commissure (AC), as illustrated by a blue vertical line on the lateral view of the 3D rendered brain image from this case. Note that the contrast between these subcortical areas is distinct in different MRI parameters. ***Abbreviations:*** CC-corpus callosum; cd-caudate nucleus; LD-lateral dorsal nucleus; LGN-lateral geniculate nucleus; MD-medial dorsal thalamic nuclei; prf-prerubral field; SN-substantia nigra; subregions of VL-ventral lateral and VP-ventral posterior nuclei. ***Sulci:*** ls-lateral sulcus; sts-superior temporal sulcus.

We then converted the delineated two-dimensional subcortical/deep brain regions into a 3D volume. We adapted this new *ex-vivo* 3D volume with 251 delineated regions as “SAM” (SAM stands for **S**ubcortical **A**tlas of **M**armoset; Figs. 4E-F, 5). This *ex-vivo* atlas was then integrated into the AFNI (Analysis of Functional NeuroImages; (Cox 1996; Saad and Reynolds 2012) and SUMA (Surface Mapper; (Cox 1996; Saad and Reynolds 2012) software packages with subcortical area labels. To preserve the contiguity of the regions, the subcortical regions were *modally smoothed* with a simple regularization procedure where each voxel was replaced with the most common voxel label in the immediate neighborhood around each voxel (27 voxels).

**Fig. 3.**
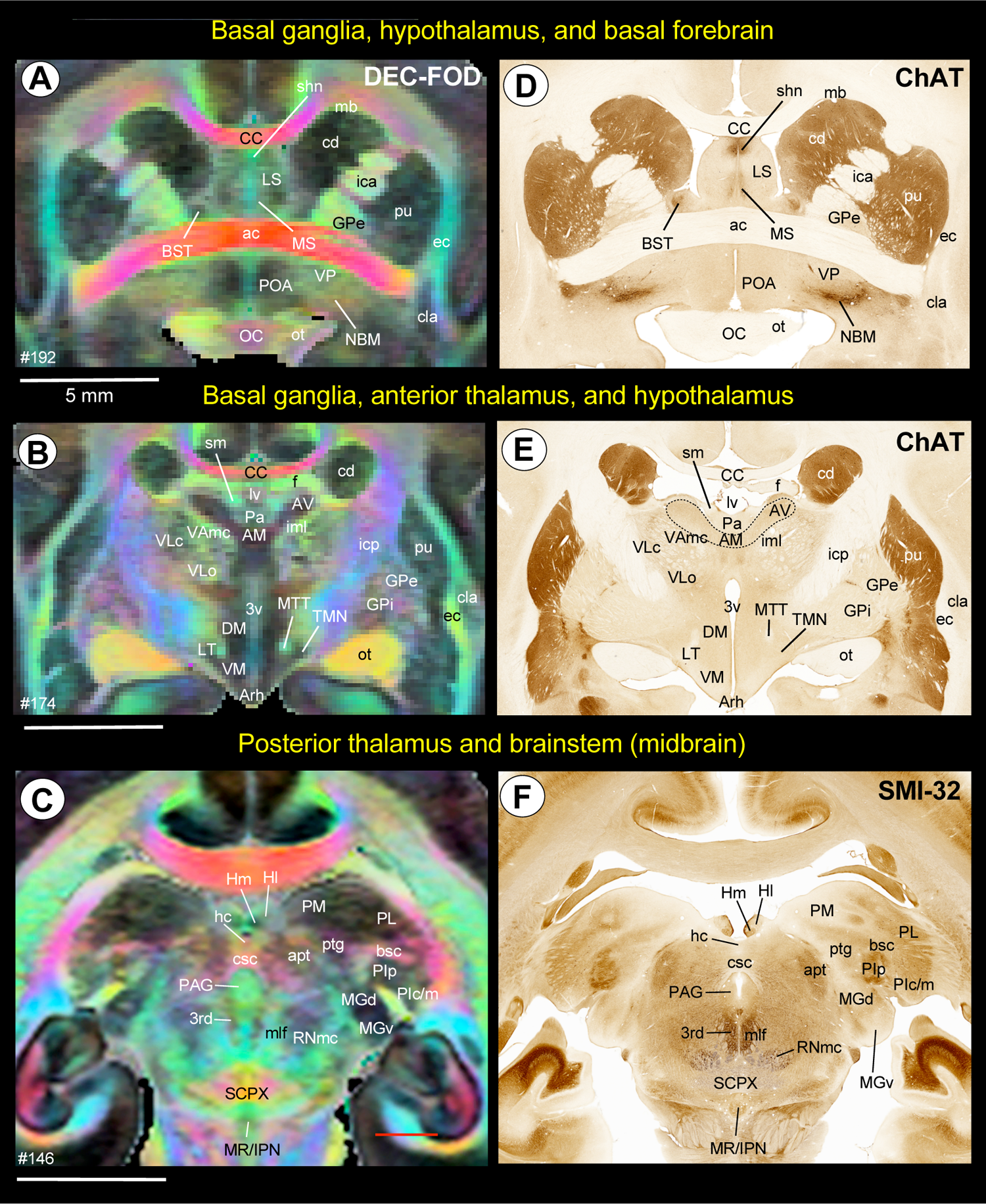
Subcortical areas for the 3D atlas (SAM). **(A-F)** Examples showing the basal ganglia, thalamus, hypothalamus, basal forebrain, and brainstem that are identified and segmented on the MAP-MRI (DEC-FOD) with reference to matched histological sections stained with ChAT and SMI-32, and other stained sections (not shown here). ***Abbreviations:*** 3^rd^-third cranial (oculomotor) nuclei; 3v-3^rd^ ventricle; ac-anterior commissure; AM-anterior medial nucleus; apt-anterior pretectal nucleus; Arh-arcuate hypothalamic nucleus; AV-anterior ventral nucleus; bsc-brachium of superior colliculus; BST-bed nucleus of stria terminals; CC-corpus callosum; cd-caudate nucleus; cla-claustrum; csc-commissure of superior colliculus; DM-dorsomedial hypothalamic area; ec-external capsule; f-fornix; GPe-globus pallidus, external segment; GPi-globus pallidus, internal segment; hc-habenular commissure; Hl-lateral habenular nucleus; Hm-medial habenular nucleus; ica-internal capsule, anterior limb; icp-internal capsule, posterior limb; iml-internal medullary lamina; IPN-interpeduncular nucleus; LS-lateral septum; LT-lateral hypothalamic area; lv-lateral ventricle; mb-Muratoff bundle; MGd-medial geniculate nucleus, dorsal division; MGv-medial geniculate nucleus, ventral division; mlf-medial longitudinal fasciculus; MR-median raphe; MS-medial septum; MTT-mammillothalamic tract; NBM-nucleus basalis of Myenert; oc-optic chiasm; ot-optic tract; Pa-paraventricular nucleus; PAG-periaqueductal gray; PIc-inferior pulvinar, caudal division; PIm-inferior pulvinar, medial division; PIp-inferior pulvinar, posterior division; PL-lateral pulvinar; PM-medial pulvinar; POA-preoptic area; ptg-posterior thalamic group; pu-putamen; RNmc-red nucleus, magnocellular division; SCPX-superior cerebellar peduncle decussation; shn-septo-hippocampal nucleus; Sm-stria medullaris; TMN-tuberomammillary nucleus; VAmc-ventral anterior nucleus, magnocellular division; VLc-ventral lateral caudal nucleus; VLo-ventral lateral oral nucleus; VM-ventromedial hypothalamic area; VP-ventral pallidum. Scale bars: 5 mm applies to A-F.

**Fig. 4.**
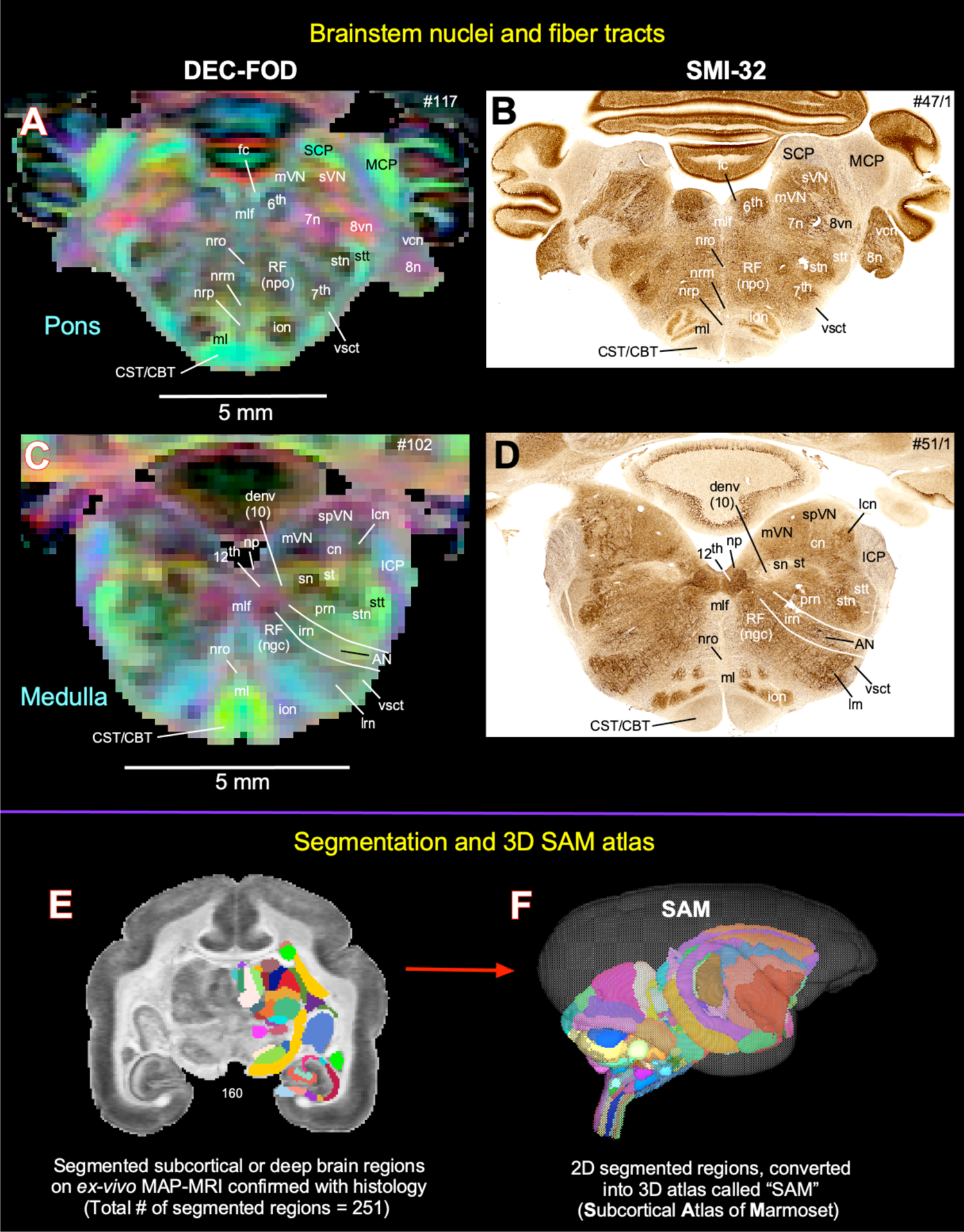
Subcortical areas for the 3D atlas (SAM). **(A-D)** More examples show the subcortical areas at the brainstem level (pons and medulla) that are identified and segmented on the MAP-MRI (DEC-FOD) with reference to matched histological sections stained with SMI-32, and other stained sections (not shown here). ***Abbreviations:*** 6^th^-abducent nuclei; 7^th^-facial nuclei; 7n-facial nerve; 8n-vestibulocochlear nerve; 8vn-vestibular nerve; 12^th^-Hypoglossal nucleus; AN-ambiguous nucleus; CBT-corticobulbar tract; cn-cuneate nucleus; CST-corticospinal tract; denv (10)-dorsal motor nucleus of vagus; fc-facial colliculus; ICP-inferior cerebellar peduncle; ion-inferior olivary nucleus; irn-intermediate reticular nucleus; lcn-lateral cuneate nucleus; lrn-lateral reticular nucleus; MCP-middle cerebellar peduncle; ml-medial lemniscus; mlf-medial longitudinal fasciculus; mVN-medial vestibular nucleus; np-nucleus prepositus; nrm-nucleus raphe magnus; nro-nucleus raphe obscurus; nrp-nucleus raphe pallidus; prn-parvicelluar reticular nucleus; RF (ngc)-reticular formation, nucleus gigantocellularis; RF (npo)-reticular formation, nucleus pontis centralis oralis; SCP-superior cerebellar peduncle; sn-solitary nucleus; spVN-spinal vestibular nucleus; st-solitary tract; stn-spinal trigeminal nucleus; stt-spinal trigeminal tract; sVN-superior vestibular nucleus; vcn-ventral cochlear nucleus; vsct-ventral spinocerebellar tract. *<u>Subcortical segmentation and 3D ex vivo digital template atlas.</u>* **(E)** Two hundred fifty-one deep brain regions, including the hippocampal formation and the cerebellum, were manually segmented through a series of 150 μm thick MAP-MRI sections using ITK-SNAP. **(F)** A 3D isosurface rendering of the individual regions within a volume rendering of the T2W dataset. This new MRI-histology-based segmented volume (called *ex vivo* “SAM”) is registered to an *in vivo* multi-subject population-based T1W MRI volume oriented to Ear-Bar-Zero stereotaxic coordinates system (Liu *et al*. 2021) or a range of *in vivo* T2W MRI volumes of marmoset monkeys with different age groups and genders. For more details, see Figures 6 and 7.

This procedure was used in our recent macaque D99 atlas V2.0 (Saleem *et al*. 2021). The method smooths edges caused by mismatches in 2D drawings applied to a 3D shape. The dataset was then subject to manual verification and correction of areal extent and architectonic borders of different subcortical areas, again aided by histology (see above). Additionally, the dataset was tested for “lost clusters” with AFNI’s @ROI_decluster. In this procedure, each region was clustered to a minimum of half the total voxels for that label. The declustered dataset was compared to the original input volume, and differences were manually corrected and reverified for lost clusters.

In order to make a symmetric template, the left-side brain T2-weighted dataset was mirrored in its x-axis (3dLRflip), and the results were combined to create a symmetric brain. The parcellated atlas was similarly mirrored. Labels were applied to the dataset for AFNI’s *whereami* functionality to show regions interactively and for command-line region selection. The labels include a short, abbreviated name and a longer descriptive name. Representative center coordinates were assigned based on the left hemisphere regions by an internal center, the voxel location inside the region that is closest to the center of mass for that region. Ten empty slices of zero values were added to the inferior, superior, and posterior regions of the dataset for general use as a target in alignment procedures. The datasets were saved in NIFTI format identified with NIFTI space code, a brief description including the atlas version. Labels and the dataset template space were saved in the AFNI extension within the NIFTI header.

#### Registration of symmetric ex-vivo SAM atlas to test subjects

To verify the usefulness and limitations of using an *ex vivo* template and atlas for *in vivo* subject marmosets, we registered this new 3D standard symmetric SAM atlas to *in vivo* T2W MRI volumes of 6 individual (control) marmoset brains of different age groups ranging from 1 to 10 and a multi-subject population-based *in vivo* T1W template oriented to Ear Bar Zero (EBZ) stereotaxic coordinates (Liu et al. 2021) using the @animal_warper program in AFNI (Jung et al. 2021). The data set was aligned to the MTR/MAP template using center-shifting, affine, and nonlinear warp transformations. The inverted transformations were combined and applied to the atlas to bring the atlas segmentation to the native space of each marmoset. The default modal smoothing was applied here to replace each voxel with the most common neighbor in the immediate 27-voxel neighborhood. For more details, see the results section and related Figure 7. The MR scanning methods to obtain T2W and T1W images of these control animals are described in the following publications (Liu *et al*. 2021; Hata et al. 2023).

## Results

### MRI markers (Fig. 2)

Using high-resolution MRI, we first identify and delineate different subcortical regions for the 3D atlas. The MAP-MRI with different microstructural parameters and other MRI parameters showed different gray and white matter contrasts outside the cerebral cortex. In particular, PA or PA/DEC-FOD (see below), T2W, and MTR (T1-like contrast) images revealed sharp boundaries and high contrast in the deep brain regions, resulting in a clear demarcation of anatomical structures such as thalamus and other brain regions (e.g., lateral geniculate nucleus-LGN, medial dorsal nucleus of thalamus-MD, prerubral field-prf, and substantia nigra-SN) (Fig. 2). We illustrated only a few MRI parameters in Figure 2, but for other MAP-MRI microstructural parameters derived from marmoset brains, see our previous study (Saleem *et al*. 2023).

Both PA/DEC-FOD and DEC-FOD referred to the same type of contrast: the propagator anisotropy (PA)-modulated direction encoded color (DEC) (Pajevic and Pierpaoli, 1999) and fiber orientation distribution (FOD) function. Similar to our previous publications (Saleem *et al*. 2021; Saleem *et al*. 2023), we have changed the wording to DEC-FOD throughout the manuscript for consistency. In each DEC-FOD voxel, the color reflects the dominant orientation of the fiber orientation, while the intensity/brightness is proportional to the propagator anisotropy (PA) in that voxel. In this way, the DEC-FOD provides a concise and compelling way to visualize the dominant fiber orientations in regions with the highest anisotropy.

#### Histological markers (***Fig. 1***)

The location, borders, and architectonic features of subcortical gray and white matter regions observed in MRI were confirmed using adjacent and matched histology sections with multiple stains derived from the same marmoset brain specimen (see STAR methods). However, we mostly relied on the DEC-FOD MR images to identify and delineate the fiber bundles of different sizes and orientations (Figs. 3, 4). The histological stains used in this study labeled different types of neuronal cells or both cell bodies and their processes in cortical and subcortical regions (See description in Fig. 2). The immuno- and histochemical stains used in this study labeled different types of neuronal cells- or both cell bodies and fiber bundles in cortical and subcortical regions. The SMI-32 antibody recognizes a non-phosphorylated epitope of neurofilament H (Sternberger and Sternberger 1983; Goldstein et al. 1987) and stains a subpopulation of pyramidal neurons and their dendritic processes in the monkey cerebral cortex (Hof and Morrison 1995; Saleem and Logothetis 2012). It is also an important marker for a vulnerable subset of pyramidal neurons in the higher cortical areas visualized in the postmortem brain of Alzheimer’s disease cases (Hof et al. 1990; Hof and Morrison 1990; Thangavel et al. 2009). The SMI-32 can detect axonal pathology in TBI brains (Johnson et al. 2016). The antibody against ChAT recognizes cholinergic neurons and has been a valuable stain for motor neurons in the monkey and human brainstem (e.g., cranial nerve nuclei), (Horn et al. 2018).

AchE is an enzyme that catalyzes the breakdown of acetylcholine and is a valuable marker for delineating different cortical areas (Carmichael and Price 1994) and major subcortical nuclei in the thalamus and brainstem (Jones 1998; Horn *et al*. 2018). The calcium-binding protein parvalbumin (PV) was thought to play an important role in intracellular calcium homeostasis. The antibody against PV has been shown previously to recognize different types of neurons in subcortical regions and a subpopulation of non-pyramidal neurons (GABAergic) in the monkey neocortex (Jones and Hendry 1989; Jones 1998; Saleem et al. 2007). The integrated multimodal approach using multiple histological stains and various MRI parameters enabled detailed noninvasive anatomical mapping and delineation of nuclei and their subregions in the deep brain regions.

#### Delineation of selected subcortical gray and white matter regions for 3D atlas (***Figs. 3 and 4***)

Using a combined *ex vivo* MRI and histology (Figs. 1, 2), we identified and delineated 211 gray matter subregions in the deep brain structures, including the basal ganglia, thalamus, hypothalamus, brainstem (midbrain, pons, and medulla), amygdala, bed nucleus of stria terminalis, and the basal forebrain. These 211 delineated areas also include the architectonically and functionally distinct non-subcortical regions, such as different lobules of the cerebellar cortex and the hippocampal formation. In addition, we also distinguished and segmented 40 fiber tracts of different orientations and sizes associated with the basal ganglia, thalamus, brainstem, and cerebellum (see Table 1). The examples in Figures 3 and 4A-D illustrate the subcortical gray and white matter regions in MAP-MRI (DEC-FOD) that are segmented with reference to matched histological sections for the 3D atlas. Although we delineated and segmented 251 gray and white matter regions to create a 3D digital template atlas, it is beyond the scope of this study to describe and illustrate all the identified areas in this report. However, we described the detailed mapping and architectonic features of many of these delineated areas in our previous publication, (Saleem *et al*. 2023). We then generated a 3D atlas from these segmented regions, and this newly segmented volume is called *ex vivo* “SAM,” or the Subcortical Atlas of the Marmoset. Figure 4F shows the lateral view of SAM with segmented subcortical regions in 3D, superimposed on the rendered brain volume from this case.

**Table 1:**
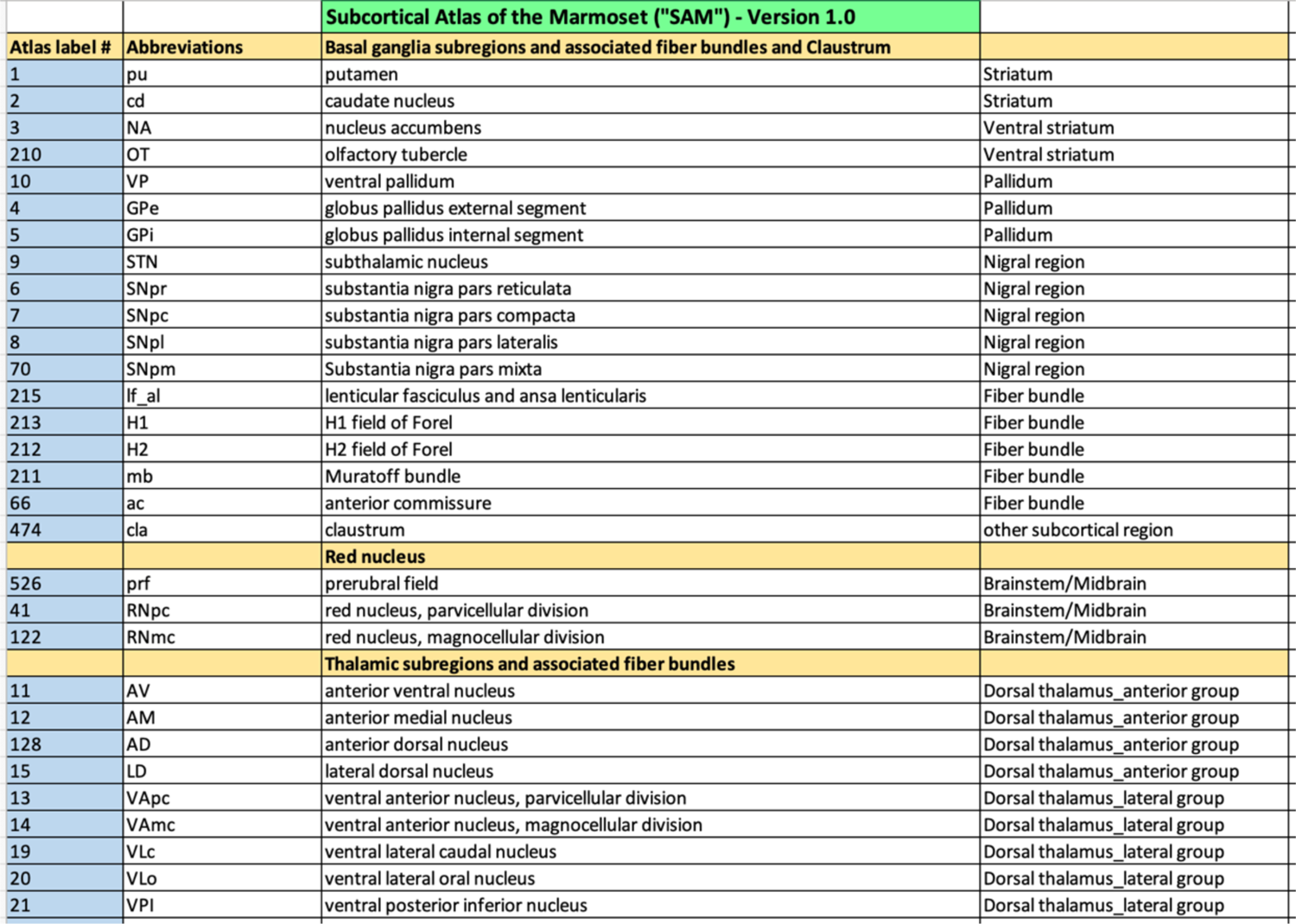
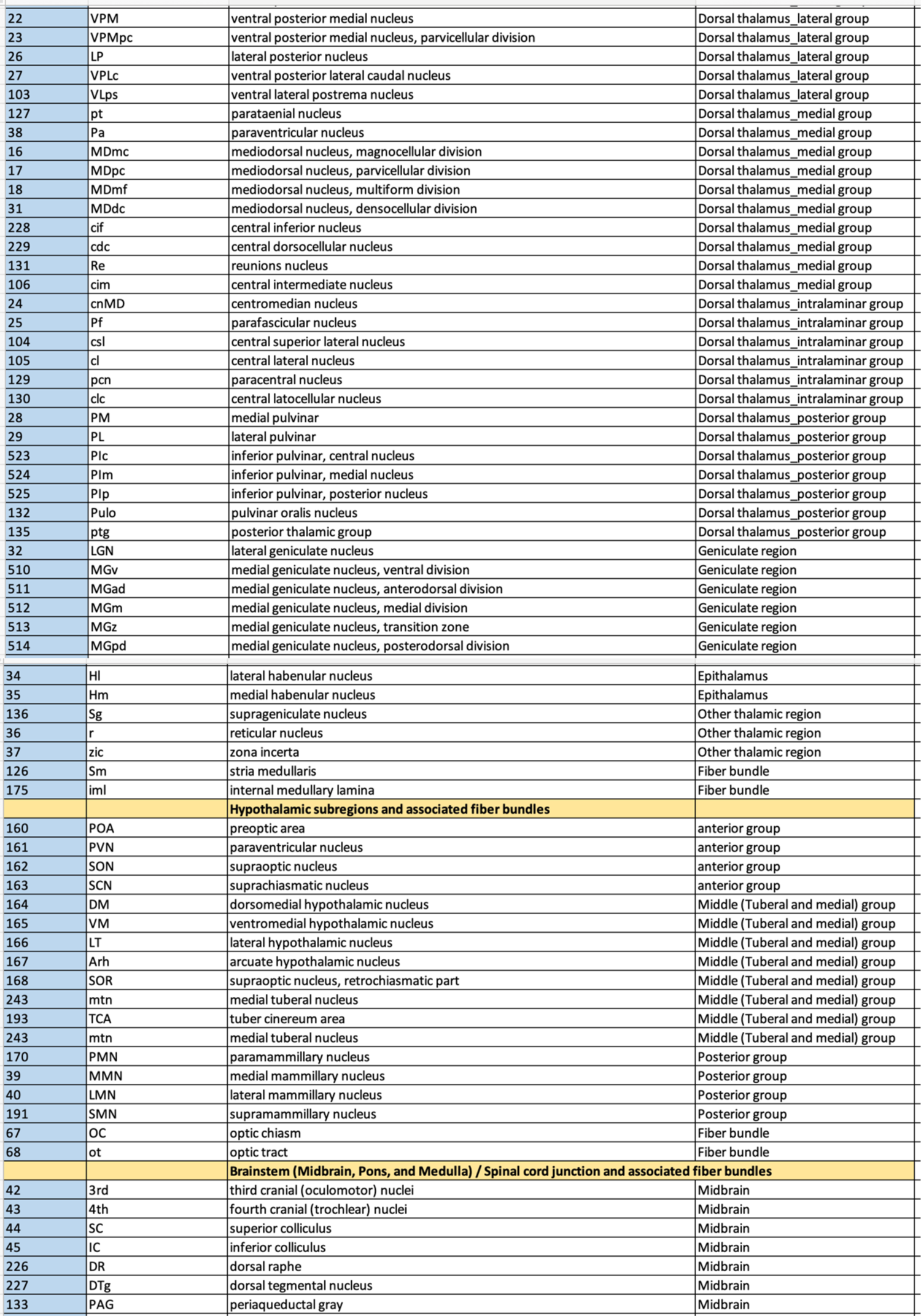
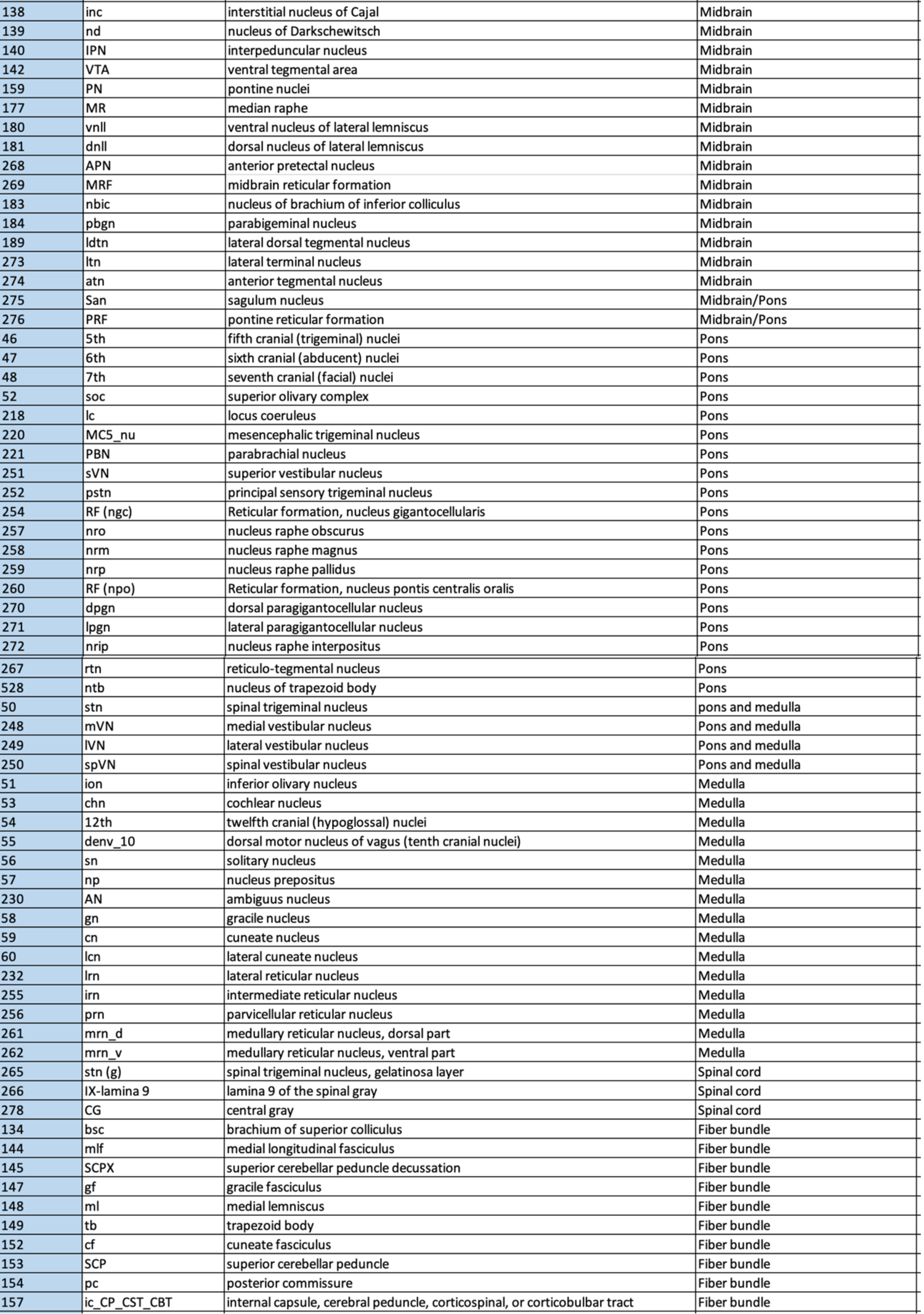
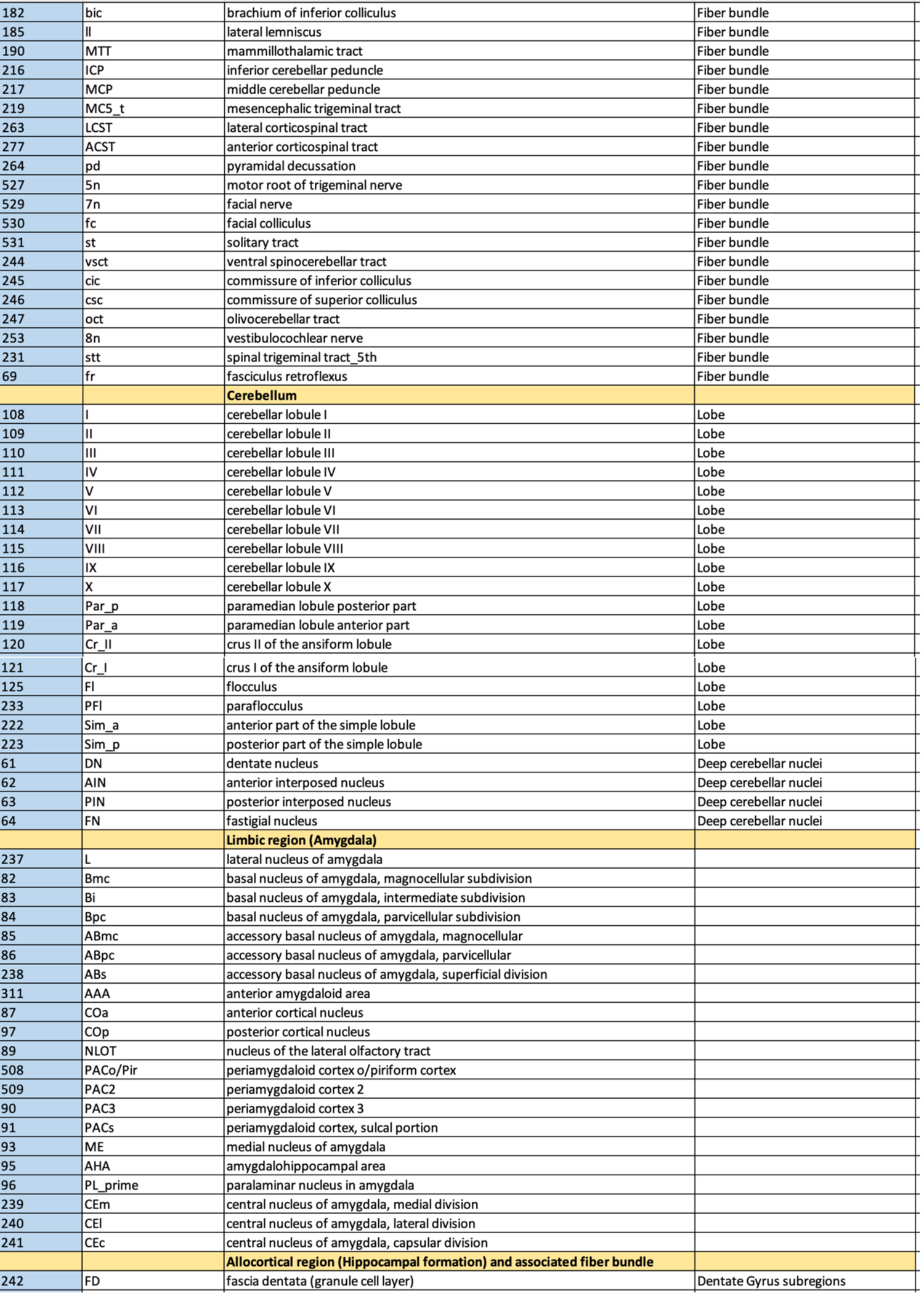
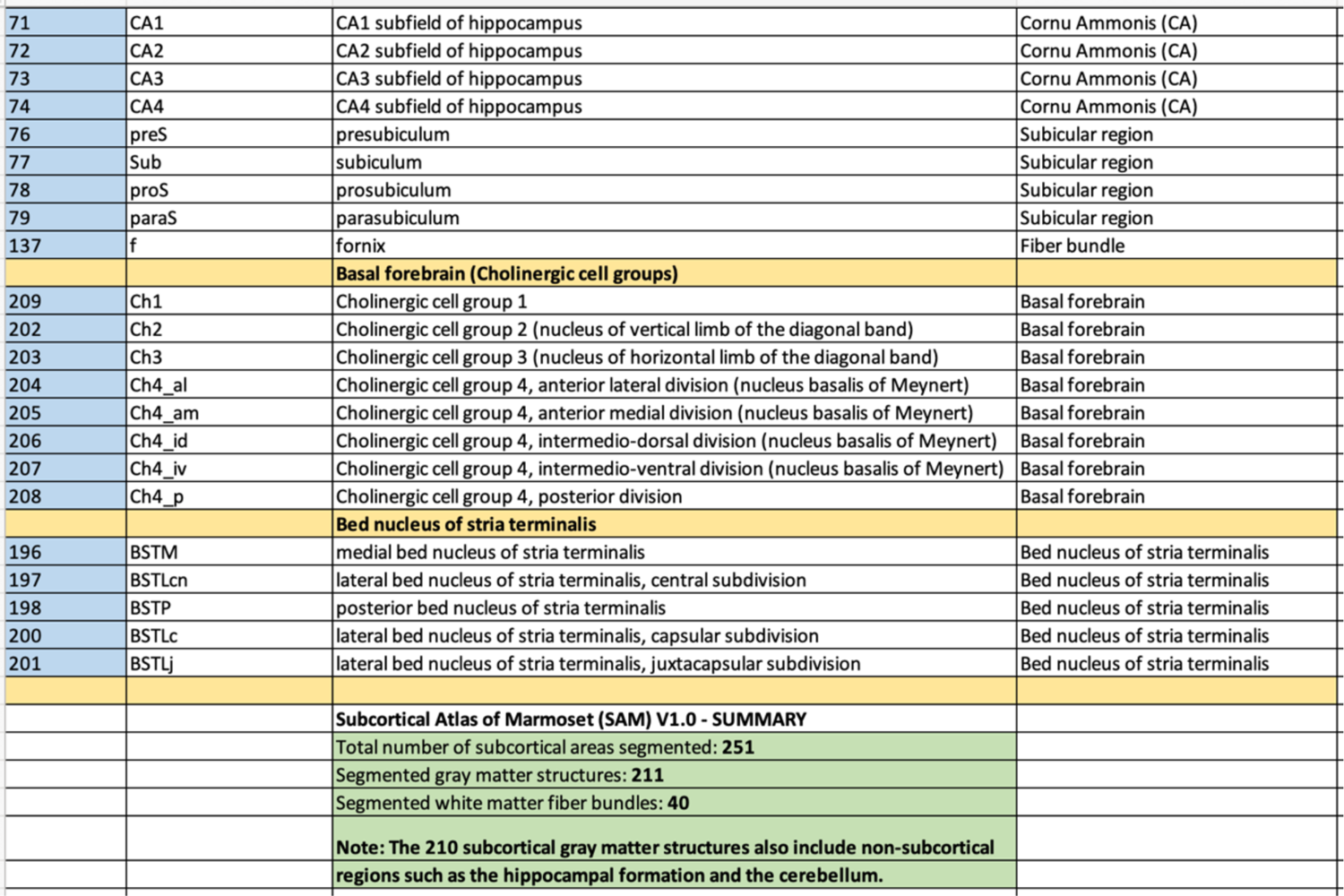

#### Ex vivo SAM template (***Fig. 5***)

Figure 5A-C illustrates the new symmetrized SAM digital template atlas with segmented subcortical regions in the horizontal, coronal, and sagittal planes of sections. We mapped 251 areas, including the subregions of the cerebellar cortex and the hippocampal formation, as described above. Figure 5D-E shows the spatial location of delineated subcortical regions on the dorsal and lateral views in 3D. This new template atlas is intended for use as a reference standard for marmoset neuroanatomical, functional, and connectional imaging studies involving subcortical targets. With AFNI’s @animal_warper, the SAM atlas can be automatically registered to the 3D anatomical scans from a wide range of marmoset subjects (see Figs. 6 and 7) and thus used to specify the areal designation relative to experimental locations of interest. This digital atlas is now available in the AFNI and SUMA analysis packages to register and apply to the brains of other individual marmoset monkeys to guide research applications for which accurate knowledge of areal boundaries is desired.

**Fig. 5.**
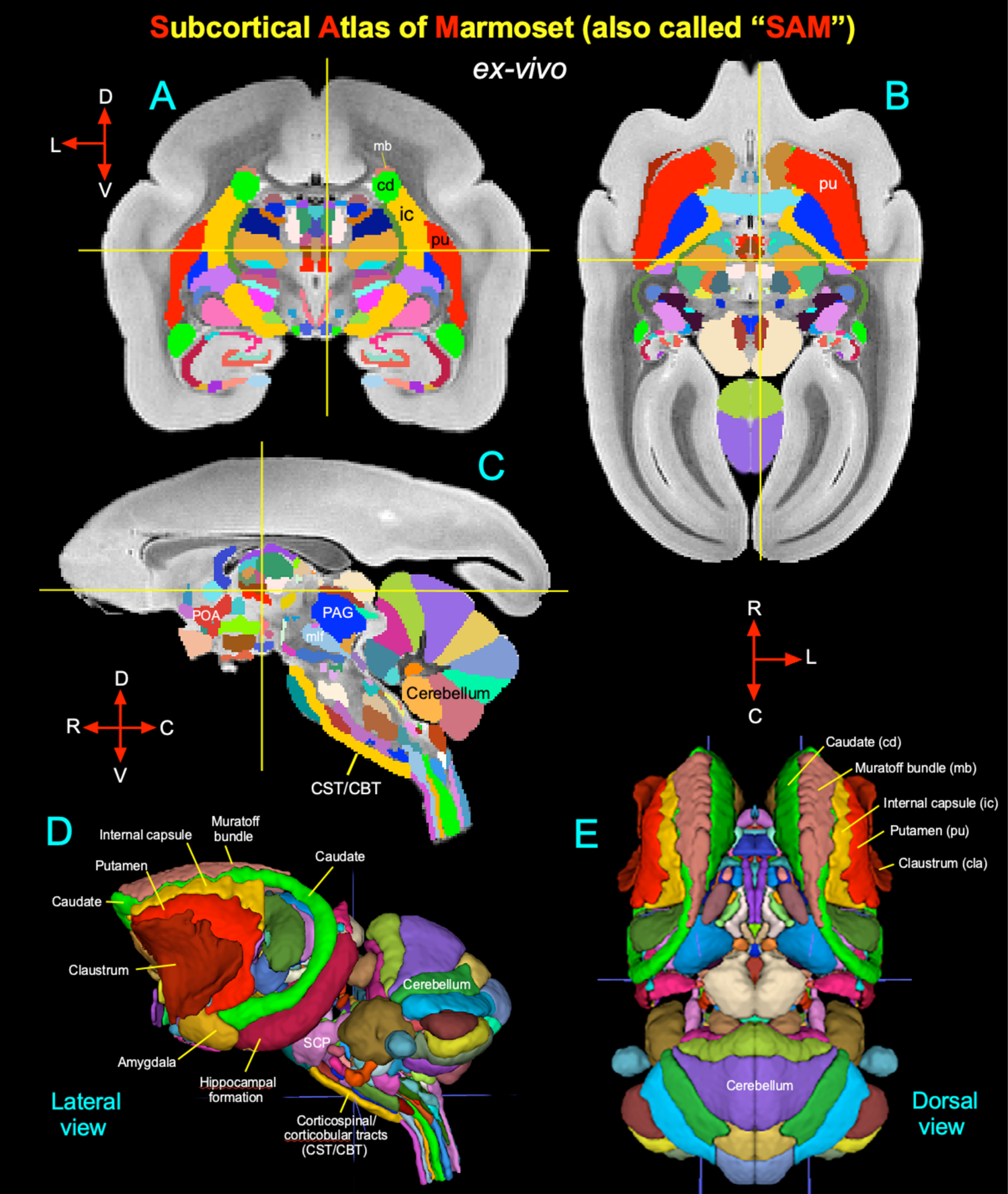
Symmetric ex vivo SAM atlas and template. **(A-C)** The “SAM” digital atlas overlaid on the coronal, horizontal, and sagittal *ex-vivo* T2W MRI template, respectively. The cross-hairs in A-C show the location of the midline thalamic subregion clc (central latocellular nucleus). **(D-E)** The spatial location of segmented subcortical regions is shown on the lateral and dorsal views in 3D. The selected subcortical regions in D-E are also indicated in A-C. ***Abbreviations:*** CBT-corticobulbar tract; CST-corticospinal tract; mlf-medial longitudinal fasciculus; PAG-periaqueductal gray; POA-preoptic area; SCP-superior cerebellar peduncle. Orientation: D-dorsal; V-ventral; R-rostral; C-caudal; L-lateral.

**Fig. 6.**
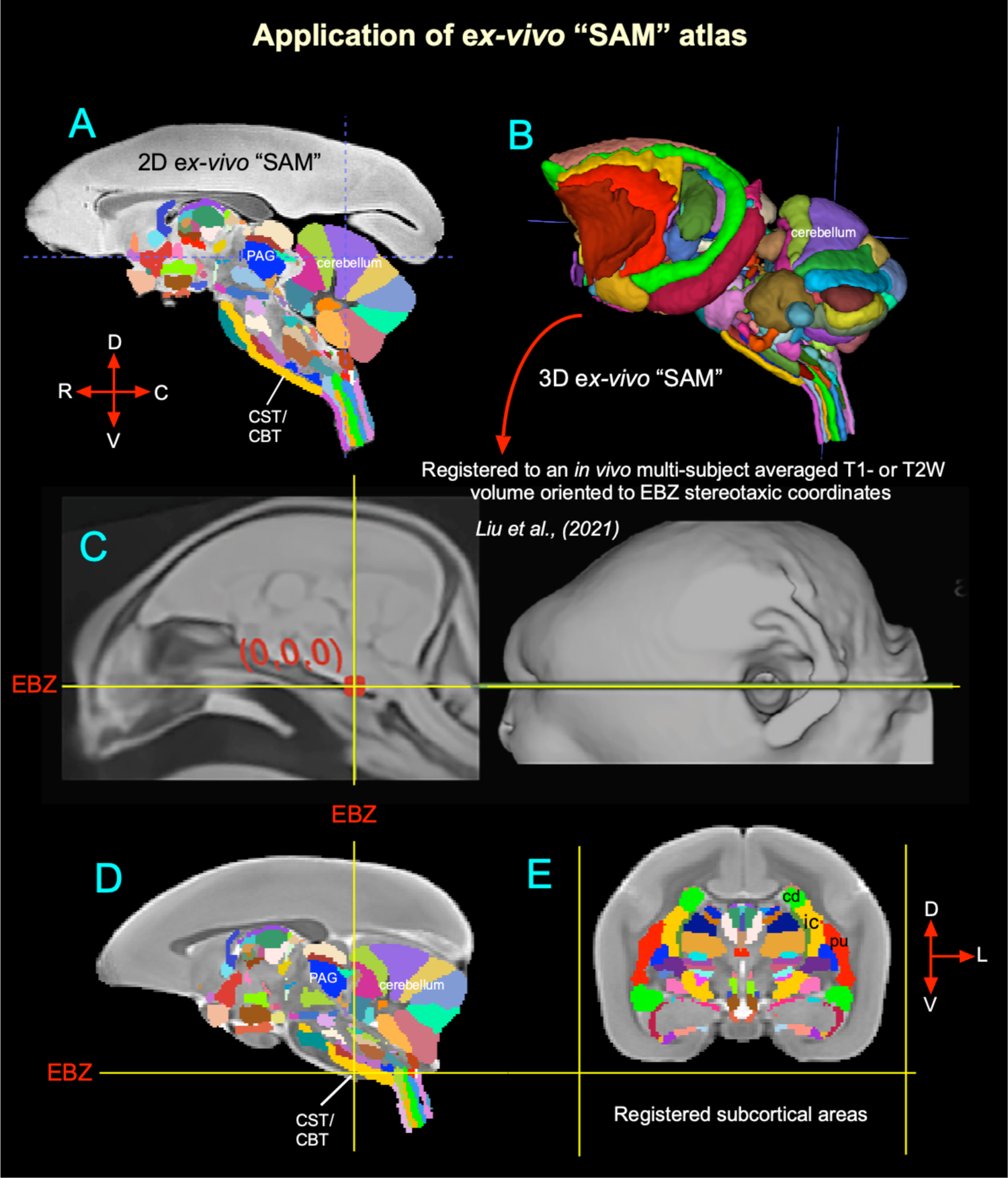
Validation of ex vivo “SAM” digital template atlas. Two hundred fifty-one deep brain regions, including the hippocampal formation and the cerebellum, were manually segmented through a series of 150 μm thick MAP-MRI or T2W images (**A**) using ITK-SNAP and derived the spatial location of these regions in 3D (**B**). This new MRI-histology-based segmented volume (called “SAM”) is registered to *in vivo* multisubject averaged T1- or T2W MRI volume (control subject) oriented to Ear Bar Zero (EBZ) stereotaxic coordinates (Liu *et al*. 2021) (**C**). The illustrations in **D-E** indicate the registered subcortical areas in this control subject (T2W). None of the registered regions in D and E were altered or adjusted. ***Abbreviations:*** CBT-corticobulbar tract; cd-caudate nucleus; CST-corticospinal tract; EBZ-ear bar zero; ic-internal capsule; PAG-periaqueductal gray; pu-putamen. Orientation: D-dorsal; V-ventral; R-rostral; C-caudal; L-lateral.

#### Application: Registering identified areas from 3D SAM to a range of test subjects (Figs. 6 and 7)

We estimated and confirmed the atlas-based areal boundaries of subcortical areas by registering this standard *ex vivo* SAM template to multiple *in vivo* T2W marmoset MRI datasets of different age groups (control adults) (Liu *et al*. 2021; Hata *et al*. 2023). To this end, we developed a novel processing pipeline within AFNI and SUMA to optimally register the new SAM atlas to *in vivo* T1W multisubject population-based (Liu *et al*. 2021) or *in vivo* T2W individual marmoset brain volumes (Figs. 6 and 7). This procedure involved a sequence of affine and nonlinear registration steps. An initial affine step gave an approximate scaling and rotation to the template. The affinely registered subject brain was gradually warped to the template by progressively smaller nonlinear warps. This procedure resulted in the subject brain data registering with the SAM template space. By inverting the combination of affine and nonlinear transformations, the atlas segmentation was warped to each subject’s original native space. The results of this pipeline are illustrated in Figures 6 and 7. In the first example (Fig. 6), the *ex vivo* SAM was registered to *in vivo* T1W multisubject averaged volume oriented to the Ear Bar Zero (EBZ) stereotaxic coordinates (Liu *et al*. 2021). Figure 6D and E show that the spatial location of transformed segmented areas (e.g., cerebellar lobules, thalamic nuclei) in this control averaged volume corresponds well with the delineated areas in the SAM atlas.

In another example in Figure 7, the corresponding location of the registered subcortical regions in the control subjects of different age groups, ranging from 1-10 years (e.g., Bmc-magnocellular subdivision of the basal nucleus in the amygdala; cla-claustrum; cd-caudate; pu-putamen; ica-anterior limb of the internal capsule) matched well with the SAM atlas (Fig. 7A-H). While determining the precise matching between the determined areas and the histologically identified regions of all six animals is a large project that is beyond the scope of the present report, these results demonstrate that a straightforward affine and nonlinear warping is sufficient to distinguish and provide atlas-based estimates of areal boundaries on marmoset subjects *in vivo*.

**Fig. 7.**
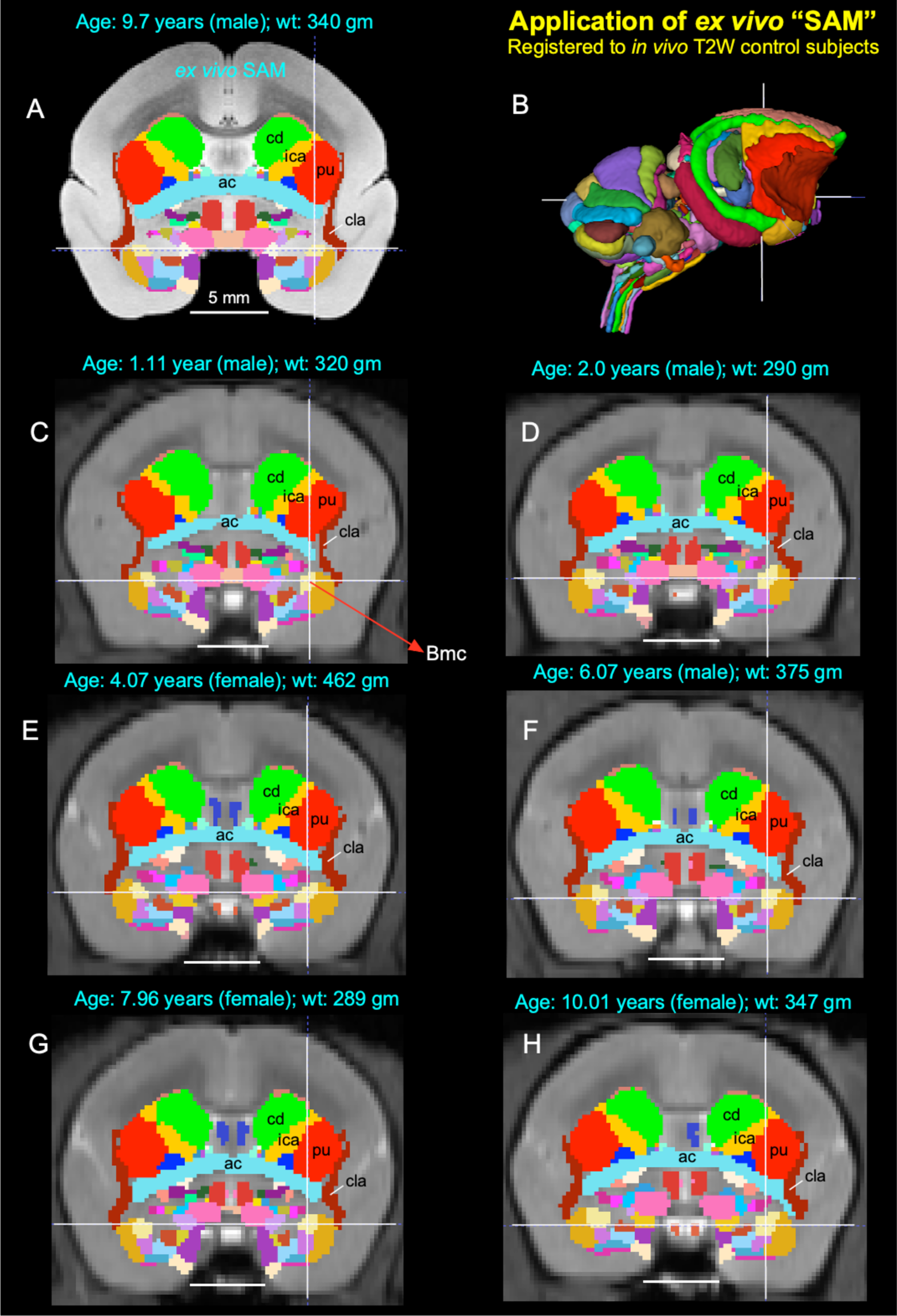
Application of 3D atlas in control subjects. Registration of SAM digital atlas to various *in vivo* T2W test subjects of different age groups, genders, and weights using a novel processing pipeline developed within AFNI (see the method section). **(A)** Mid-coronal section from SAM atlas with delineated subcortical regions. **(C-H)** Coronal slices from 6 control animals, with the SAM atlas registered to the T2W MRI volume of each animal in its native space. None of the registered regions were altered or adjusted in these animals. Note the corresponding location of the deep brain regions in the control subjects (e.g., ac-anterior commissure; Bmc-basal nucleus of the amygdala, magnocellular division, indicated by cross-hair; cd-caudate; cla-claustrum; ica-anterior limb of the internal capsule; pu-putamen) closely matched with the SAM **(A)**. The MRI volumes of these six control subjects were obtained from the publicly shared multimodal brain MRI database covering the marmosets with a wide age range (Hata *et al*. 2023). Scale bars in A-H = 5 mm.

Atlases and templates are available as both volumes and surfaces in standard NIFTI and GIFTI formats. While this 3D digital atlas can be used in other image registration and analysis software packages, here we use the AFNI and SUMA programs with their advanced atlas features for purposes of demonstration (Cox 1996; Saad and Reynolds 2012). The atlas is integrated into the most recent versions of AFNI and SUMA, making for straightforward identification of areal identity in any marmoset subject registered to the template and for the individual marmoset subject in its native space by the inverse transformations. The 3D template volume, atlas, and script for atlas registration of *in vivo* scans are now available for download through the AFNI and SUMA websites at https://afni.nimh.nih.gov/pub/dist/atlases/marmoset/SAM_Marmoset/SAM_marmoset_subcortical_dist.tgz The AFNI software can install this simply with the @Install_SAM_Marmoset command.

## Discussion

Despite its essential role as a research model for human brain development and neurological disorders, the marmoset monkey lacked a comprehensive, well-organized MRI-histology-based atlas of subcortical regions. In this study, we have generated a 3D digital template atlas of the marmoset from 251 segmented subcortical regions (called “SAM”) using high-resolution MAP-MRI, T2W, and MTR images, combined and correlated with the histological sections of the same brain specimen. Our results demonstrate that, at a high spatial resolution, the combined use of MRI parameters and matched histology sections with five different stains enabled detailed noninvasive segmentation of gray and white matter regions in the deep brain structures. This integrated multimodal approach yields a more objective and reproducible delineation of nuclei and their boundaries in the deep brain structures, which include the basal ganglia, thalamus, hypothalamus, limbic region (amygdala), basal forebrain, and the rostrocaudal extent of the brainstem (midbrain, pons, and medulla). Many of these deep brain targets and their subregions are less prominent or indistinguishable from neighboring structures with conventional T1W or T2W MRI volumes (Saleem *et al*. 2023). This new atlas is intended for use as a reference standard template for neuroanatomical, functional (fMRI), clinical, and connectional imaging studies involving subcortical targets in marmoset monkeys. We also estimated and confirmed the atlas-based areal boundaries of subcortical areas by registering this *ex vivo* atlas template to *in vivo* T1- or T2W MRI datasets of marmoset control adults (single and multisubject population-based volumes), using a novel pipeline developed within AFNI. In the following sections, we compare our new marmoset digital atlas, “SAM,” with other available atlases in the field and highlight some advantages of the present offering.

### Standard SAM versus other marmosets MRI- and histology-based atlases

The present marmoset digital template atlas of subcortical regions, derived from the MRI and histology, is one of the few digital atlases created in recent years. In one study, Liu and colleagues (Liu *et al*. 2018) constructed a 3D digital atlas of the marmoset brain based on MR image contrasts observed in *ex vivo* MTR, T2W, and diffusion MR images. This study manually delineated 54 cortical but only 16 subcortical areas in their digital atlas. It also lacks the parcellation of brainstem structures (midbrain, pons, and medulla) and nuclei within the major subcortical structures like the thalamus and the amygdala (for example). No histology information is available from this study. Although this MRI-based atlas is helpful for some applications, delineating the cytoarchitectonic areas based on MRI contrasts alone, without corresponding and matched histological information from the same brain specimen to serve as a control, may produce inaccurate boundaries that include additional gray and white matter regions leading to biases in size and volume estimation of ROI. As indicated in our previous macaque (Saleem *et al*. 2021) and marmoset (Saleem *et al*. 2023) studies, the multimodal MRI parameters acquired with high-spatial resolution (100-200 μm), aided by histology derived from the same brain specimen, is key to delineating nuclei and their subregions for the construction of 3D digital template atlases. In another study, Majka and colleagues (Majka *et al*. 2020; Majka *et al*. 2021) created a Nencki-Monash template, a probabilistic atlas based on the morphological average of 20 young marmoset brains obtained by 3D reconstructions generated from Nissl stained serial sections. It provided a cytoarchitectonic parcellation of cortical areas but no subcortical or deep brain regions.

In a different study, Hashikawa and colleagues (Hashikawa *et al*. 2015) reconstructed a series of Nissl-stained axial slices into a 3D brain model with cortical and subcortical parcellations using a volume-rendering method. They also reproduced virtual low-resolution parasagittal and coronal slices from this axially generated 3D volume. The introduction of the histology-based template atlases and space was an important step forward and offered a good estimate of areal boundaries and virtual brain structures delineated by their histological features in 3D space. While useful for some applications, this approach does not attempt to preserve the native geometry of the brain due to the distortion and shrinkage of histological sections during section cutting, staining, and mounting. Thus, the spatial accuracy of the resulting volumetric or surface atlas remains questionable. The chemoarchitectonic characterization and parcellation are also necessary for delineating several brain structures in cortical and subcortical regions (Saleem et al., 2021, 2023), but these features are lacking in these studies.

The present digital atlas avoids geometrical transformations of the cytoarchitectonic information from histological sections to match the layout of the animal’s brain. Such transformations can introduce registration errors that are difficult to correct. The most important unique feature of our subcortical atlas (SAM) is the strict adherence to an MRI scan with the adjacent and matched histology sections with multiple histo- and immunohistochemical stains from the same brain. As a result, the alignment accuracy between areal boundaries and gross anatomical features is optimized for identifying regions of interest in the present study (Figs. 4 and 5 in this study; 4-9 in Saleem et al., 2013).

### Generalization and Validation of 3D Atlas

The *ex vivo* SAM volume is registered to multiple *in vivo* 3D templates of different age groups using widely available tools of whole-brain MRI registration. When applied to the 3D volume, the transformation derived from this warping allows for the labeling of subcortical targets in the brains of individual animals as accurately as possible. It also integrates the information directly with their anatomical and functional imaging results in surface modes. The compilation of any brain atlas, which includes the assignment of boundaries and names to individual areas, is an inherently imperfect endeavor whose main goal is to provide a common anatomical framework for a range of research projects and data. In the present case, the innovation rests on the creation of a 3D digital marmoset atlas whose anatomical borders were, from the outset, created based on MR-registered histological sections. This digital atlas (SAM) is based on the precise histological borders from one particular monkey, which, because of the initial registration to the MRI from the same animal, can be represented on the brain of any experimental animal via an alignment procedure such as the one used in this study.

A complete validation of this 3D atlas, such as estimating the architectonic boundaries between different areas for a population of marmoset brains, is beyond the scope of the present report. Nonetheless, our multipronged analysis supports the validity of the SAM template by estimating the architectonic boundaries between different subcortical/deep brain areas for a population of *in vivo* T1- or T2W marmoset brains. In this analysis, we revealed that the MRI registration procedure using SAM (Figs. 6, 7) could be smoothly applied to test subjects of different genders, age groups (1-10 years old), and sizes (290-462 gm). Thus, it is possible to estimate the histological boundaries of subcortical areas in any marmoset monkey (Fig. 7). The validation of brain regions in a given subject is helpful for neurosurgical navigation of electrode or implantable devices to a potential target for deep brain stimulation (DBS) in non-human primate models of psychiatric or neurological disorders (e.g., Parkinson’s disease (Min et al. 2016; Vitek and Johnson 2019). It is also useful for localizing labeled neurons and terminals after anatomical tracer injections, fMRI activation regions (Baker et al. 2006; Logothetis et al. 2012; Turchi et al. 2018; Murris et al. 2020), and mapping the trajectories of subcortical development from young to adult marmoset monkeys (Seki et al. 2017; Sawiak et al. 2018).

### MAP-MRI-based atlases in humans (Future directions)

In this study, we delineated and generated a 3D subcortical atlas of the marmoset monkey (SAM) using high-resolution MAP-MRI and other MRI parameters and matched histological sections with multiple stains derived from the same brain specimen. We further illustrated how our atlas is used to locate small subcortical structures after registration to T1- or T2W MRI volumes of control subjects acquired *in vivo*. These results suggest the utility of a high-resolution atlas in studies of marmoset monkey disease models and highlight the potential of high-resolution MAP-MRI in delineating small subcortical structures based on differences in microstructural properties. The spatial resolution of MAP-MRI data acquired on clinical scanners is significantly lower, approximately 1-2 mm, than what was used here. Nevertheless, promising advances in the gradient coil design (McNab et al. 2013), RF coil engineering (Keil et al. 2013; Truong et al. 2014), dMRI pulse sequence design (Avram, Guidon, et al. 2014), and spatial encoding (Feinberg et al. 2010; Setsompop et al. 2018) are expected to significantly improve spatial resolution and SNR to allow submillimeter clinical MAP-MRI scans in the near future (Huang et al. 2021). Concurrently, new clinically feasible diffusion encoding strategies (Avram et al. 2010; Avram et al. 2019; Avram et al. 2021) and analyses (Avram, Saleem and Basser 2022; Magdoom et al. 2023) are being developed to characterize and delineate specific microscopic tissue water pools without the need to increase the spatial resolutions. Taken together, these advances will enable the construction of high-resolution cortical and subcortical maps and atlas templates of the human brain that will improve localization of neurosurgical navigation, fMRI responses, high-precision placement of recording and stimulating electrodes in patients with Parkinson’s, epilepsy, mild TBI (mTBI), and other diseases.

### Summary and Conclusion

We have created a comprehensive MRI template and corresponding digital atlas of subcortical regions in the marmoset brain based upon a large set of histological and very high-resolution structural MRI and MAP-MRI data for a single marmoset. The atlas provides a usable standard for region definition, while the template provides a standard reference and space. This standard space allows for marmoset research to be reported on a common basis across research sites and across marmoset monkeys. As used in human studies with MNI or Talairach space, this target template space provides a platform for voxelwise group analysis. Additionally, the atlas allows for automated analysis against a set of standard region locations, either in the template space or in the native space of the individual subjects. The current atlas, template MRI data sets, surfaces, and user scripts for aligning individual subjects to this template are publicly available at the following link: https://afni.nimh.nih.gov/pub/dist/atlases/marmoset/SAM_Marmoset/SAM_marmoset_subcortical_dist.tgz

## Key Resources Table

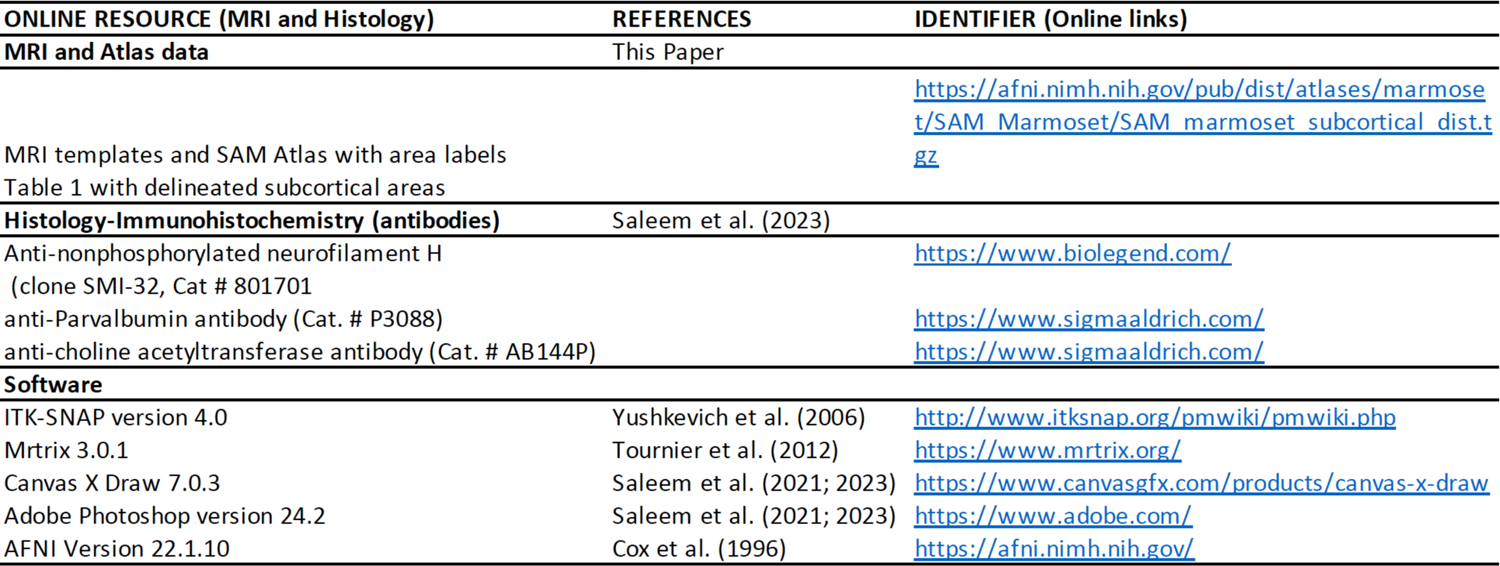

## Credit author statement

***Kadharbatcha S. Saleem:*** Corresponding author, designed the study, coordinated the project, prepared the specimen for MRI and histology; mapped, segmented, and verified all the anatomical regions of interest with reference to MRI and histology; generated new subcortical 3D atlas template/Table, made illustrations; wrote, edited, and streamlined the manuscript.

***Alexandru V Avram:*** Conducted all MRI experiments, including MAP-MRI scans, processed and analyzed all MRI data, helped with atlas data registration, and wrote and edited the manuscript.

***Daniel Glen:*** Integrated the atlas dataset into AFNI and SUMA software packages, wrote code for the atlas region regularization, helped with atlas data registration, and edited the manuscript.

***Vincent Schram:*** Obtained high-resolution images of all histology-stained sections in the microscope imaging center (MIC) at NICHD and commented on the manuscript.

***Peter J Basser:*** helped design the study, edited the manuscript, and provided research resources.

## Acknowledgments

This work was supported by the CNRM Neuroradiology/Neuropathology Correlation/Integration Core, 309698-4.01-65310 (CNRM-89-9921); Intramural Research Program of the *Eunice Kennedy Shriver* National Institute of Child Health and Human Development; the Intramural Research Program of the National Institute of Neurological Disorders and Stroke; “Connectome 1.0: Developing the next generation human MRI scanner for bridging studies of the micro-, meso -and macro-connectome”, NIH BRAIN Initiative 1U01EB026996-01. We thank James Pickel, Transgenic core facility at NIMH, for providing perfusion-fixed marmoset monkey brain for our experiments; Cecil Chern-Chyi Yen for helping with the initial setup for the MRI scanning; and Michal Komlosh for the preparation of the specimen for MRI. Finally, we thank the Microscope imaging core (MIC) at NICHD for help with the high-resolution imaging of histology sections. All histological processing of the brain tissue was done by Dr. Du and his team at FD NeuroTechnologies in Columbia, Maryland.

